# A Survey of Orbitrap All Ion Fragmentation Analysis Assessed by an R MetaboList Package to Study Small-molecule Metabolites

**DOI:** 10.1101/257147

**Authors:** Enrique Sentandreu, Manuel D Peris-Díaz, Shannon R Sweeney, Jennifer Chiou, Nathalie Muñoz, Stefano Tiziani

**Author notes:** Correspondence to whom should be addressed (S.T.). These authors contributed equally to the manuscript (E.S., M.D.P-D., S.R.S.).

## Abstract

Leukemia cell and melanoma tumor tissue extracts were studied for small (mostly *m/z* <250) polar metabolites by LC-ESI-HRMS^n^ analysis powered by a hybrid Quadrupole-Orbitrap. MS data was simultaneously acquired in fast polarity switching mode operating in MS^1^ and MS/MS (All Ion Fragmentation, AIF) full-scan analyses at high mass resolution. Positive assignments were achieved by AIF analysis considering at least two characteristic transitions of metabolites. A targeted metabolite profiling was achieved by the relative quantification of 18 metabolites through spiking their respective deuterated counterparts. Manual data processing of MS^1^ and AIF scans were compared for accurate determination of natural metabolites and their deuterated analogs by chromatographic alignment and peak area integration. Evaluation of manual and automated (MetaboList R package) AIF data processing yielded comparable results. The versatility of AIF analysis also enabled the untargeted metabolite profiling of leukemia and melanoma samples in which 22 and 53 compounds were respectively identified outside those studied by labeling. The main limitation of the method was that low abundance metabolites with scan rates below 8 scans/peak could not be accurately quantified by AIF analysis. Combination of AIF analysis with MetaboList R package represents an opportunity to move towards automated, faster and more global metabolomics approaches supported by an entirely flexible open source automated data processing platform freely available from Comprehensive R Archive Network (CRAN, https://CRAN.R-project.org/package=MetaboList).

## INTRODUCTION

Metabolomics, the youngest of the “-omics” disciplines has presented more analytical challenges than its predecessors, genomics and proteomics, due to the broad physical and chemical properties of metabolites [1-4]. In its truest form, metabolomics is completely untargeted [1]. However, this presents several challenges for data acquisition and data processing [2,5,6]. Often a compromise must be made between the aim to measure the entire metabolome with the reality of resources available. Specifically, the election between targeted and untargeted approaches in metabolomics research is commonly ruled by the analytical technology available at the time of analysis [7]. In the case of mass analyzers, commonly coupled to chromatographic techniques, electronics and device hardware play an essential role in choosing the metabolomics approach to follow. Ideally, metabolomics-oriented platforms of analysis should exhibit high sensitivity, versatility, robustness, scan rate, and mass accuracy, with a special dedication to the qualitative/quantitative analysis of small molecules [8]. Moreover, metabolomics workflows must include a reliable automated processing of the large data sets generated by this type of analysis [2,9]. As a result, the current degree of complexity achieved by mass spectrometry-based metabolite studies has and continues to promote the development of versatile, simple, and high-throughput methodologies that can facilitate the activity of researchers [7,10,11].

A hybrid Quadrupole-Orbitrap liquid chromatography (LC) coupled mass analyzer is a versatile analytical solution considering its high sensitivity, mass accuracy, scan speed, and dynamic range-duty cycle [11]. The all ion fragmentation (AIF) technology, which applies a Higher-energy Collisional Dissociation (HCD) fragmentation to all ionized molecules without mass filtering (quadrupole not engaged), is one of many different operating modes for this instrumental setup. AIF is a data independent analysis (DIA) that was first introduced in early orbitrap detectors as a full-scan MS/MS operation mode that permits the acquisition of high mass-accuracy fragmentation data of all metabolites in a complex mixture by time. In this approach, multiple data-dependent MS/MS (dd-MS^2^) parameters such as inclusion/exclusion lists of precursors, Top N precursors, inclusion/exclusion times, and number of MS/MS scans per analyte, are not applied. Such complexity in the dd-MS^2^ experimental design can limit the extension of qualitative analysis, especially for low abundance metabolites not present in inclusion and Top N lists. In addition, quantitative analysis is not possible when operating in dd-MS^2^ mode because it is not a full-scan experiment. The combination of MS^1^ and AIF full-scans provides an opportunity to perform retrospective data analysis of additional compounds of interest based on hypotheses that arise later [11,12], thus providing dimensions of flexibility not achieved by other MS^2^ analyses such as multiple reaction monitoring (MRM) [11]. As pointed out by Bateman *et al.* (2009) [13], AIF analysis can provide improved mass accuracy and requires negligible time investment to develop methods in contrast to that required for traditional QQQ-based MRM analysis. Advances in Time of Flight (TOF) instrumentation, standalone or coupled to a quadrupole (Q-TOF), has also resulted in instruments capable of operating at high scan rate/mass resolution in a way similar to AIF conditions (i.e. All Ions MS/MS, MS^E^, and MS^ALL^). However, efficiency in the study of small molecules with *m/z* <300 by TOF analysis is often hindered by technical limitations such as lower mass accuracy, dynamic range, and multiplexing performance compared to an orbitrap, so the application of MetaboList with TOF instruments should be evaluated separately [8,13-15]. Nonetheless, researchers using these technologies may also take advantage of the data analysis workflow utilizing the automated R package MetaboList evaluated here.

Currently, AIF analysis has been mainly used in untargeted lipidomics studies where the ability to uncover lipid-specific fragments allows validation of lipid species present [12,16-18]. The versatility of AIF beyond lipidomics has been explored by identification of some pharmaceutical metabolites [19] and isotopically labeled polar and non-polar metabolites [20], indicating the potential for widespread application provided an efficient data processing method is available. As mentioned above, data generated by AIF is the resultant of a full-scan MS/MS analysis that pools HCD fragments from all ionized molecules over time. However, automated processing of AIF data cannot be performed in the same way as full scan MS^1^ (intact molecules) since the generated raw data file is in MS^2^ format but without precursor ions, which are required for traditional data/time dependent analyses including most traditional MS/MS processing programs. Manual or semi-automated AIF data processing has resulted in largely targeted/non-extensive methodology commonly used for quantitative-qualitative metabolomics [13,21] and proteomics [22] research, mainly for the purpose of validating a limited number of ambiguous assignments [23,24]. Among the scarce alternatives that are currently available to carry out the automated processing of LC-AIF data, two freely available options MS-DIAL [25] and MetDIA [26], should be highlighted for their efficiency. The former was originally inspired by Gas Chromatography-Mass Spectrometry (GC-MS) deconvolution processing whereas the latter is partially supported by the R software environment. Although MS-DIAL was successfully tested in lipidomics research, its lower efficiency for small molecule analysis was demonstrated by [26], suggesting the usefulness of a more customizable framework for data processing in contrast to the rigid online algorithms of analysis currently available. It should be noted that both data processing solutions were assayed using a LC-Q-TOF device with mass tolerances beyond 15 ppm and XCMS-based peak picking processing, so there may be opportunities for improvement in data processing with lower mass tolerances and/or alternative peak picking methods [26].

To date, all of the options available to carry out qualitative analysis of AIF data have relied on classical spectral matching. As such, breakdown patterns from isolated metabolites (precursors) are compared with those from their respective AIF counterparts. Unfortunately, AIF breakdown often induces over-fragmentation (further fragmentation of already-formed fragments) of molecules [22], thus relative abundances of AIF fragments can be rather far from those achieved through the isolation and subsequent fragmentation of precursors (i.e. MRM). As a result of this limitation, the question regarding the efficiency of an entirely customizable solution that does not require classical spectral matching or peak picking (such as XCMS or other commercially available software) for automated processing of AIF data still remains unclear. In this line, considering the results from Li *et al.* [26] and the efficiency previously demonstrated by R programming packages for the automated processing of MS^1^ data [9,27], we can conclude that this is an important challenge to overcome in order to maximize the usefulness of AIF analysis. This work aims to demonstrate the utility of combining high-resolution MS^1^ and MS/MS AIF full-scan analyses with entirely automated data processing powered by the newly developed R-package MetaboList as an easy, versatile, affordable, and reliable workflow for global metabolomics [28]. The methodology proposed here represents an independent, but complementary, alternative to current strategies that rely on expensive, commercially available software and/or other freely available R-based packages that utilize spectral matching approaches for automation of AIF data processing.

Strengths and limitations of high-resolution MS^1^ and MS/MS AIF full-scan analyses in conjunction with automated R package MetaboList processing are discussed in this study to demonstrate usefulness and flexibility of this methodology regarding metabolomics research. The small (mainly at *m/z* < 250) polar metabolites detected in leukemia cell and melanoma tumor tissue extracts were exhaustively investigated. Furthermore, this work aims to set up the workflow analysis for the easy implementation of the developed methodology in mass spectrometry-based metabolomics with independence of the sample source.

## MATERIALS AND METHODS

### Chemicals and materials

LC-MS grade formic acid (FA), acetonitrile (ACN) methanol (MeOH), dimethyl sulfoxide (DMSO), and ammonium formate (AF) were from Fisher Scientific (Pittsburgh, PA, USA). Water was of ultrapure grade (EMD Millipore Co., Billerica, MA, USA). Deuterated standards D4-Anthranilic acid, D5-Kynurenic acid, D4-Kynurenine, D3-Quinolinic acid, and D3-3-Hydroxy-DL-kynurenine were purchased from Buchem BV (Apeldoorn, The Netherlands). Deuterated standards D2-Fumaric acid, D3-DL-Glutamic acid, D3-Malic acid, D4-Citric acid, D4-succinic acid, D2-Cysteine, D4-Alanine, D2-Glycine, D5-Glutamine, D3-Serine, D3-Aspartic acid, D4-Cystine, and D5-L-Tryptophan were purchased from Cambridge Isotope Laboratories Inc. (Tewksbury, MA, USA). Stable isotopically internal standards were in the range of 98-99% chemical purity and their isotopic purity was in the 97-99% range with the exception of D5-Kynurenic acid which main isotopologue (45% of the appeared D1-D5 cluster signal) corresponded to the D3-form. Commercial negative/positive calibration solutions for the MS device were from Thermo Fisher Scientific (San José, CA, USA).

### Internal standards

Targeted quantitative analysis was performed by spiking samples with deuterated standards. Two different batches of internal standard mixtures were prepared according the sample analyzed. The leukemia cell extract was spiked with all the deuterated standards available at the time of this research which were fumaric acid, glutamic acid, malic acid, citric acid, succinic acid, cysteine, alanine, glycine, glutamine, serine, aspartic acid, cystine and tryptophan standards all previously dissolved in water with 0.2% FA (MIX 1). Analysis of tumor tissue was oriented toward the study of the kynurenine cycle and were spiked with deuterated anthranilic acid, kynurenic acid, kynurenine, quinolinic acid, 3-hydroxy-kynurenine, and tryptophan standards dissolved in MeOH/water (50:50) with 2% DMSO (MIX 2).

### Sample preparation

Leukemia cells (Pediatric T-cell Acute Lymphoblastic Leukemia derived from a primary patient sample received from the Dell Children’s Blood & Cancer Center, Austin, TX) were cultured under standard conditions, at 5% CO_2_ and 37°C, in RPMI 1640 medium supplemented with 10% fetal bovine serum (FBS; Hyclone Laboratories, Logan, UT, USA). Cells and medium were collected and gently centrifuged at 1400 rpm for 5 minutes. Medium supernatant was aspirated and the cell pellet was washed twice with cold phosphate buffered solution (PBS; Hyclone Laboratories, Logan, UT, USA). Cells were transferred to an Eppendorf tube and pulse centrifuged to pellet. The supernatant was aspirated and the cell pellet was snap frozen in liquid nitrogen and stored at −80°C until extraction. A modified Bligh-Dyer method for metabolite extraction was used [29]. Briefly, the cell pellet was resuspended in 1 mL chilled water/MeOH (50:50) and transferred to a 2 mL glass vial containing 0.5 mL cold chloroform. The glass vial was vortexed on a platform shaker for 10 minutes at 2500 rpm and then centrifuged for 20 minutes at 4750 rpm at 4°C to achieve phase separation. The polar phase was removed, transferred to an Eppendorf tube, and dried in a CentriVap Concentrator (Labconco, Kansas City, MO, USA) at 4°C. The dried polar phase was resuspended in 200 μL of ultrapure water containing 0.2 ppm of a deuterated internal standard mixture (MIX 1). Insoluble particulates were removed from the sample by ultrafiltration with a washed Nanosep 3K Omega centrifugal filter (Pall Corporation, Port Washington, NY, USA) at 8000 rpm and 4°C for 20 minutes [30]. The filtrate was transferred into a glass LC-MS vial and stored at −80°C until injection.

The well-established murine model of human melanoma, B16-OVA (purchased from ATCC, Manassas, VA, USA), was allografted into wild type C57BL/6J mice [31]. Procedures were approved by the Institutional Animal Care and Use Committee (IACUC) at The University of Texas at Austin prior to any murine experiments. Mice were euthanized by CO_2_ asphyxiation and cervical dislocation at a maximal tumor size of 200 mm^2^ and before to any signs of distress were detected. Postmortem, 100 mg tumor tissue aliquot was transferred to a 2 mL tissue homogenization tube with mixed beads and 0.5 mL of chilled water/MeOH (50:50) was added. The sample was homogenized in a Precellys-24 cryo homogenizer (Bertin Technologies, Saint Quentin en Yvelines, France) at −5°C and 5000 rpm for two cycles of 20 seconds each. The lysate was recovered and transferred into a glass vial containing 0.5 mL of chilled chloroform. The homogenization tube was washed with an additional 0.5 mL of chilled water/MeOH (50:50) and added to the glass vial for metabolite extraction. The glass vial was vortexed at 1500 rpm for 3 minutes, centrifuged at 4750rpm for 20 minutes at 4°C, and the methanolic supernatant was transferred to an Eppendorf tube and dried in a CentriVap vacuum concentrator. The sample was resuspended in 150 μL of ultrapure water containing 0.2 ppm of a deuterated internal standard mixture (MIX 2), filtered, and stored as described above.

### Liquid chromatography high-resolution mass spectrometry analysis (LC-HRMS)

Chromatographic analysis was performed on an Accela HPLC system equipped with a quaternary pump, vacuum degasser and an open autosampler with a temperature controller (Fisher Scientific, San José, CA, USA). Chromatographic separation of metabolites was achieved by hydrophilic interaction liquid chromatography (HILIC) and reverse phase (RP) approaches. HILIC analysis was performed on a ZIC p-HILIC 100×2.1 mm, 5μm particle size column (Millipore Co., Billerica, MA, USA) with the following analytical conditions: solvent A, water/FA (99.9:0.1) containing 10 mM AF; solvent B, ACN/FA (99.9:0.1); separation gradient, initially 96% B, linear 96-20% B in 15 minutes, purging with 1% B for 5 minutes and column equilibration with 96% B for 10 minutes; flow rate, 0.3 mL/min; injection volume, 1.5 μL. RP analysis was conducted on a 150 mm×2.1 mm, 3 μm particle size Synergi-Hydro C18 column (Phenomenex Inc, Torrance, CA, USA) with the following separation conditions: solvent A, water/FA (99.8:0.2); solvent B, ACN; separation gradient, initially 1% B, held for 2 minutes and then linear 30-80% B in 8 minutes, washing with 98% B for 5 minutes and column equilibration with 1% B for 15 minutes; flow rate, 0.25mL/min; injection volume, 3μL. In all cases, autosampler and column temperatures were set at 6°C and 22°C, respectively.

Mass spectrometry analysis was carried out on a Q Exactive Hybrid Quadrupole-Orbitrap benchtop detector equipped with an electrospray (ESI) source simultaneously operating in fast negative/positive ion switching mode (Thermo Scientific, Bremen, Germany). Multiplexing capabilities of the analyzer led to combine full-scan MS^1^ (full MS) and full-scan MS/MS (AIF) experiments with settings: microscans, 1; AGC target, 1e^6^; maximum injection time, 100 ms; mass resolution, 35000 FWHM at *m/z* 200 for full MS analysis whereas AIF scan conditions were: microscans, 1; AGC target, 3e^6^; maximum injection time, 1000 ms; mass resolution, 70000 FWHM at *m/z* 200; HCD energy, 30. Larger AGC and maximum injection time values in AIF scan aimed to preserve sensitivity compromised by over-fragmentation of HCD ions. In both cases, the instrument was set to spray voltage, 4.0 kV; capillary temperature, 300°C; sheath gas, 55 (arbitrary units); auxiliary gas, 30 (arbitrary units); *m/z* range, 50-650; data acquisition, centroid mode.

A complementary targeted MS/MS analysis was carried out to obtain the characteristic breakdown pattern of 3-hydroxykynurenine since no HCD information in positive ionization mode was provided by the mzCloud database (https://www.mzcloud.org, HighChem LLC, Slovakia). A merged full MS and dd-MS^2^ analysis was performed in the melanoma sample using the same analytical conditions mentioned above with some modifications: MS/MS inclusion list for masses at *m/z* 225.0868 and 228.1058 (3-hydroxykynurenine and D3-hydroxykynurenine, respectively); MS/MS AGC target and injection time of 1e^6^ and 100 ms, respectively; number of MS/MS scans, 3. It must be highlighted that in this research, the term MS/MS is indistinctly used throughout the text for both AIF and dd-MS^2^ analyses since both generate breakdown ions. Thereby, differentiation among full-scan and data-dependent fragmentation analyses is done according to their abbreviations (AIF and dd-MS^2^, respectively).

Accuracy of MS analysis was ensured by calibrating the detector using the commercial calibration solutions provided by the manufacturer followed by a customized adjustment for small molecular masses. Masses at *m/z* 87.00877 (Pyruvic acid); 117.01624 (D2-Fumaric acid); 149.06471 (D3-Glutamic acid); 265.14790 (Sodium dodecyl sulfate) and 514.288441 (Sodium taurocholate) were used for the negative ionization mode whereas masses at *m/z* 74.09643 (n-Butylamine), 138.06619 (Caffeine fragment), 195.08765 (Caffeine) and 524.26496 (Met-Arg-Phe-Ala tetrapeptide, MRFA) were used to adjust mass accuracy of the positive ionization mode. Mass tolerance was kept at 5 ppm in both full-scan MS and AIF modes. The LC-MS platform of analysis was controlled by a PC operating the Xcalibur v. 2.2 SP1.48 software package (Thermo Scientific, San Jose, CA, USA).

### LC-MS data analysis

Samples were studied combining targeted and untargeted approaches. Since samples were spiked with labelled standards, targeted qualitative and quantitative analysis was carried out by mimicking MRM experiments considering two characteristic transitions from the examined metabolite. Spiked standards in this analysis assisted the identification of their respective natural counterparts through the appropriate alignment of the molecular and respective AIF ions. Customized MetaboList MS^1^ and MS^2^ libraries ([M+H]^+^ and [M-H]^-^ in.csv format) listing the labelled standards and their respective natural metabolites were built for manual and automated data processing for targeted analysis considering a mass tolerance of 5 ppm. As a rule, two main fragments detailed in the mzCloud database at HCD of 30 for the natural metabolites were considered as well as their respective labelled counterparts, using the most abundant as the quantitative ion. To avoid interferences caused by considering targeted fragments with very small molecular masses, commonly shared with other coeluted species in AIF analysis, other major fragments with higher molecular masses were chosen as an alternative (i.e. kynurenine in melanoma sample). Relative quantitative analysis was based on peak area ratios between quantitative ions belonging to natural metabolites in samples and their respective deuterated counterparts. Initially, quantitative analysis was manually performed using Xcalibur to compare results from full MS and AIF ratios. Next, manual and automated AIF results were compared to evaluate the robustness of the R-based MetaboList package. No biological replicates were considered since accuracy of the automated quantification was evaluated by using the values from manual analysis of the same sample to avoid discrepancies caused by sampling deviations.

Untargeted analysis of samples aimed to investigate the qualitative performance of the automated data processing. It was more complex than the targeted strategy described above since all of the characteristic MS/MS fragments that can ensure a positive assignment were considered. The untargeted analysis was divided into two steps. A preliminary metabolite profiling of samples was carried out through the study of full-scan MS^1^ data from both positive and negative ionization modes using the Thermo SIEVE v 2.2.58 SP2 program (Thermo Fisher Sci., San José, CA, USA). An in-house library (.csv format) listing the neutral molecular mass of 300 small (m/z <650) polar metabolites commonly found in biological analyses was interrogated, considering a mass tolerance of 5 ppm as an initial discriminant constraint. Initial positive assignments were subsequently confirmed by automated processing through the comparison of their AIF breakdown fragments with those described in the mzCloud database for each respective precursor ion and ionization mode. To achieve this goal, a customized MetaboList library listing the characteristic MS/MS data of the preliminary SIEVE assignments was built ([M+H]^+^ and [M-H]^-^ in.csv format) to be loaded during the automated processing of AIF data. Since AIF analysis induces over-fragmentation of ions and this phenomenon is also influenced by the abundance of the metabolites in samples, this untargeted MS^2^ library was built considering fragments detailed in mzCloud database in an HCD range of 30-50 and with a minimum relative abundance of 20%.

Evidently, proper alignment of intact (SIEVE analysis) and AIF (MetaboList analysis) ions led to achieve positive identifications. Identical MS^1^ results were found by MetaboList automated processing of data loading the aforementioned SIEVE library but in [M+H]^+^ and [M-H]^-^ input format. Possibility of loading customized libraries by MetaboList analysis listing neutral mass of candidates is currently in progress.

### Automated data processing

To facilitate the understanding of how the R-based strategy addressed concerns in automated data processing, **Fig. 1** summarizes the workflow proposed in this study. Prior to R processing, original LC-MS data files (.raw extension) were converted to .mzXML files through MSconvert from Proteowizard (http://proteowizard.sourceforge.net) to separate full MS and AIF experiments with each scan still merging positive/negative analyses [32].

**Figure 1.**
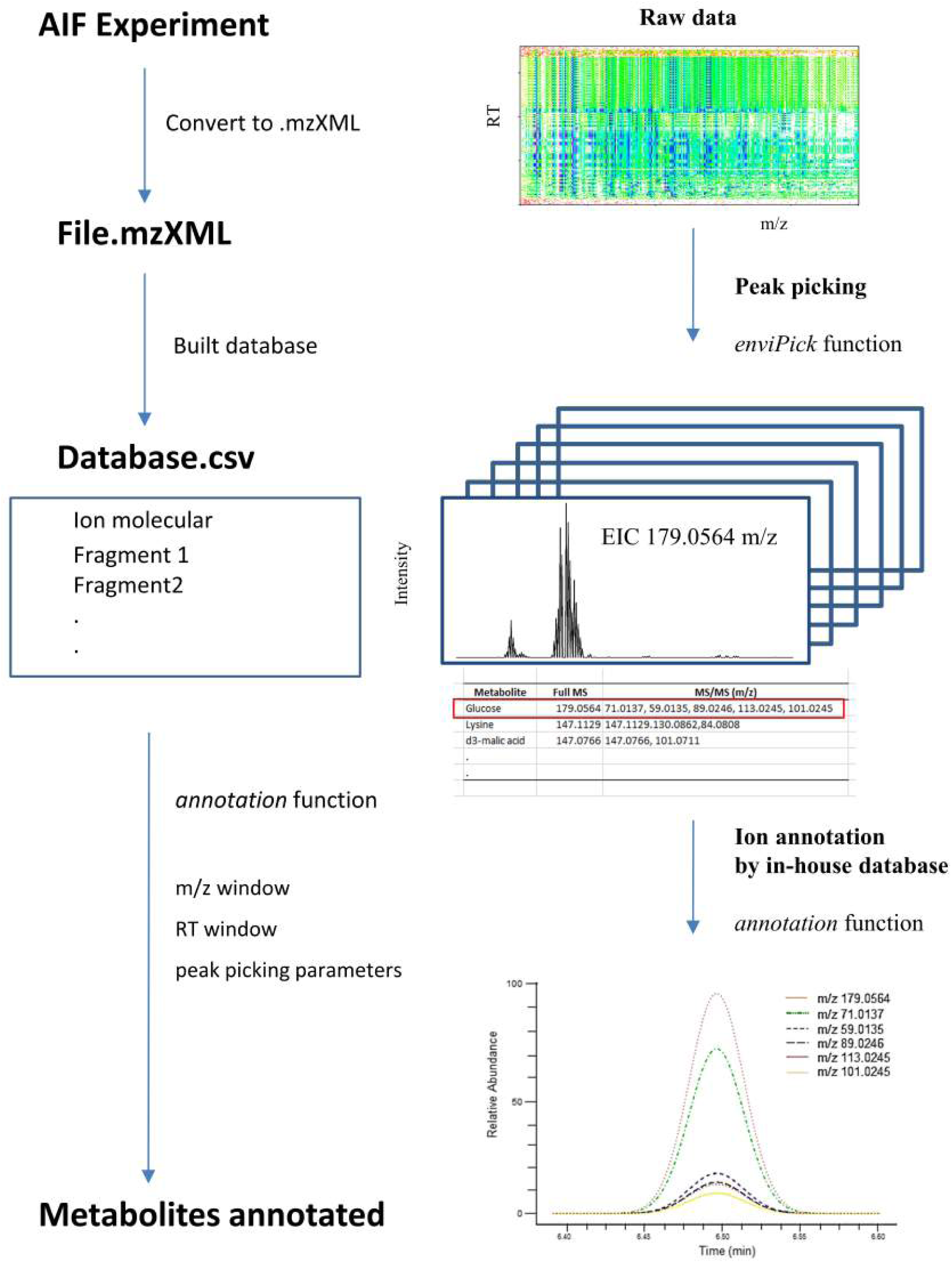
Automated R workflow analysis proposed in this research.

AIF scans were handled by the R package MetaboList [28] available at the Comprehensive R Archive Network (CRAN) repository (https://cran.r-project.org/web/packages/MetaboList). Peak picking was conducted in three consecutive steps, by incorporating the R package enviPick [33] embedded in the R package MetaboList. Initially, an agglomerative partitioning approach was performed of the respective positive/negative full-scan MS/MS data into individual partitions through retention time (rt) and *m/z* gap widths (settings at drtgap = 25-1000 and dmzgap = 1-5, respectively). Secondly, clustering of extracted ion chromatograms (EICs) was carried out in the subsets of the partitions generated (drtdens = 2 and dmzdens = 5). Finally, peak picking of EIC clusters given a retention time window (drtsmall = 5-20), was restricted to the ranges: minint =1e5, maxint= 1e9, SB = 0.1-4 and SN = 0.01-2. Further details on peak picking performed are described in the enviPick manual [33].

In addition, MetaboList supports automated ion annotation by loading the aforementioned untargeted hybrid MS-MS/MS library with the characteristic intact masses and respective breakdown patterns of the preliminary SIEVE analysis. The algorithm generates an EIC matrix and searches for ions defined in the database, thus creating a new subset with only those ions with similar *m/z* values according to a mass tolerance of 5 ppm. Then, the subset was filtered by comparing the retention times of intact and respective fragment masses according to a specified time deviation of 4 seconds to ensure appropriate alignment of ions for positive assignment. A faster device and/or reduction of the number of scan events could further decrease time deviations. It should be noted that retention times have only been used in this study as a constraint for time deviations among aligned fragments belonging to the same compound. However, for well-known compounds it can also be used as a discriminant parameter for targeted analysis by its inclusion into a customized library.

## RESULTS AND DISCUSSION

### Instrumental Parameters

Assayed MS conditions of analysis had an average scan rate of 2.7 scans/second for the entire full MS/AIF duty cycle; that is, 0.7 scan/second per event (four events considering polarity switching). This rate requires chromatographic peaks widths of around 0.3 minutes to obtain the recommended 10-12 scans per metabolite peak in MRM analysis and avoided the use of Ultra High Pressure Liquid Chromatography (UHPLC) conditions. A reduction in scan modes (operating exclusively in AIF mode) and/or using a faster device would facilitate the analysis of narrower peaks.

### Targeted analysis: Full MS vs. AIF Analysis

Initially, manual data processing was performed by Xcalibur to validate capacity of AIF analysis for qualitative/quantitative determination compared to full MS study. **Fig. 2** shows peak representation of intact MS and AIF scans for the serine/D3-serine pair in leukemia sample. Very clearly, a positive assignment was achieved by the alignment of the molecular masses and quantitative AIF fragments detailed in **Table 1**. Chromatographic and MS properties of target metabolites and deuterated standards in leukemia and melanoma samples and their respective full-scan MS and AIF ratios from peak area integration are listed in **Table 1**. MS^1^ and AIF ratios from manual integration were rather comparable (deviations around 15%, **Table 1**) in most cases indicating the usefulness of AIF for quantitative analysis. Discrepancies observed were the consequence of lower natural metabolite abundance that was directly translated into low scan/peak values. As an example, serine achieved full MS and AIF rates of 13 and 12 scan/peak with ratios of 0.1992 and 0.1907 (deviation of 4.46%), respectively. In contrast, the kynurenic/D3-kynurenic acid pair in the melanoma sample, even with a good MS^1^ and AIF peak alignment (same as showed in **Fig. 2** for serine), had rather different ratios of 0.0387 and 0.0606 (deviation of 36%), respectively. The intensity of natural kynurenic acid in the sample was 1x10^5^ which resulted in scan rates of 8 and 6 scans/peak for full MS and AIF analyses respectively, thus the subsequent misquantification. Similarly, conflicting results were obtained for cystine in the leukemia sample with an AIF scan rate of only 5 scans/peak. The most extreme case of intensity dependence was found for quinolinic acid in the melanoma sample with MS^1^ and AIF peaks defined by 10 and 1 scans/peak, respectively. This outstanding peak rate difference is most likely the result of strong over-fragmentation of the AIF quantitative ion. Despite deviations in the quantification analysis of low-abundant metabolites, AIF approach was effective at generating qualitative data that enabled positive assignments.

**Figure 2.**
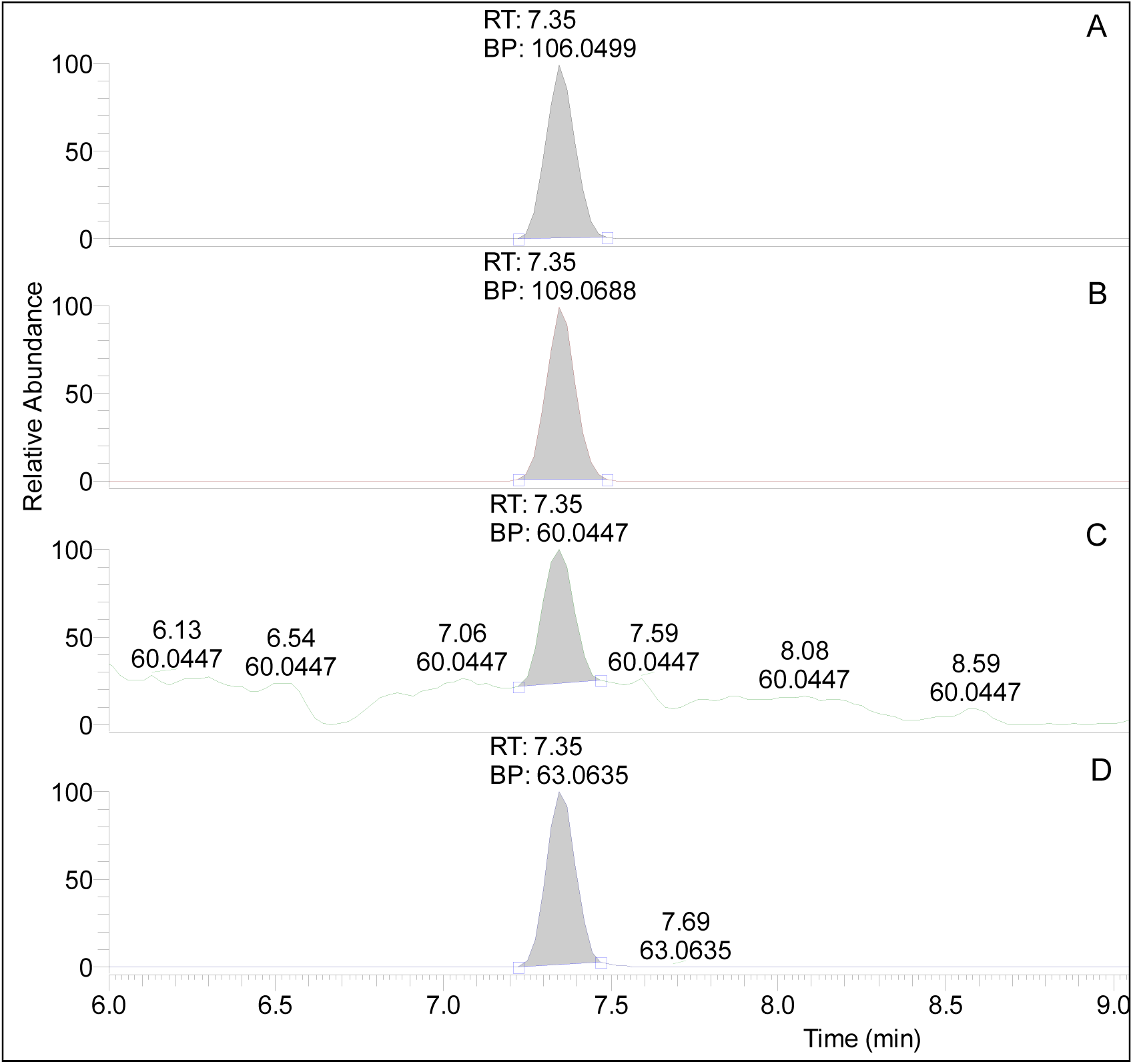
Full MS and AIF scan representation in positive ionization mode of serine/D3-serine pair in leukemia sample: A. Molecular mass of serine, B. Molecular mass of D3-serine, C. AIF quantitative fragment of serine, D. AIF quantitative fragment of D3-serine. Peak area integration from manual data processing indicated in gray. Nomenclature used: RT, chromatographic retention time; BP, base peak corresponding to intact and MS/MS ions (Table 1).

**Table 1.**
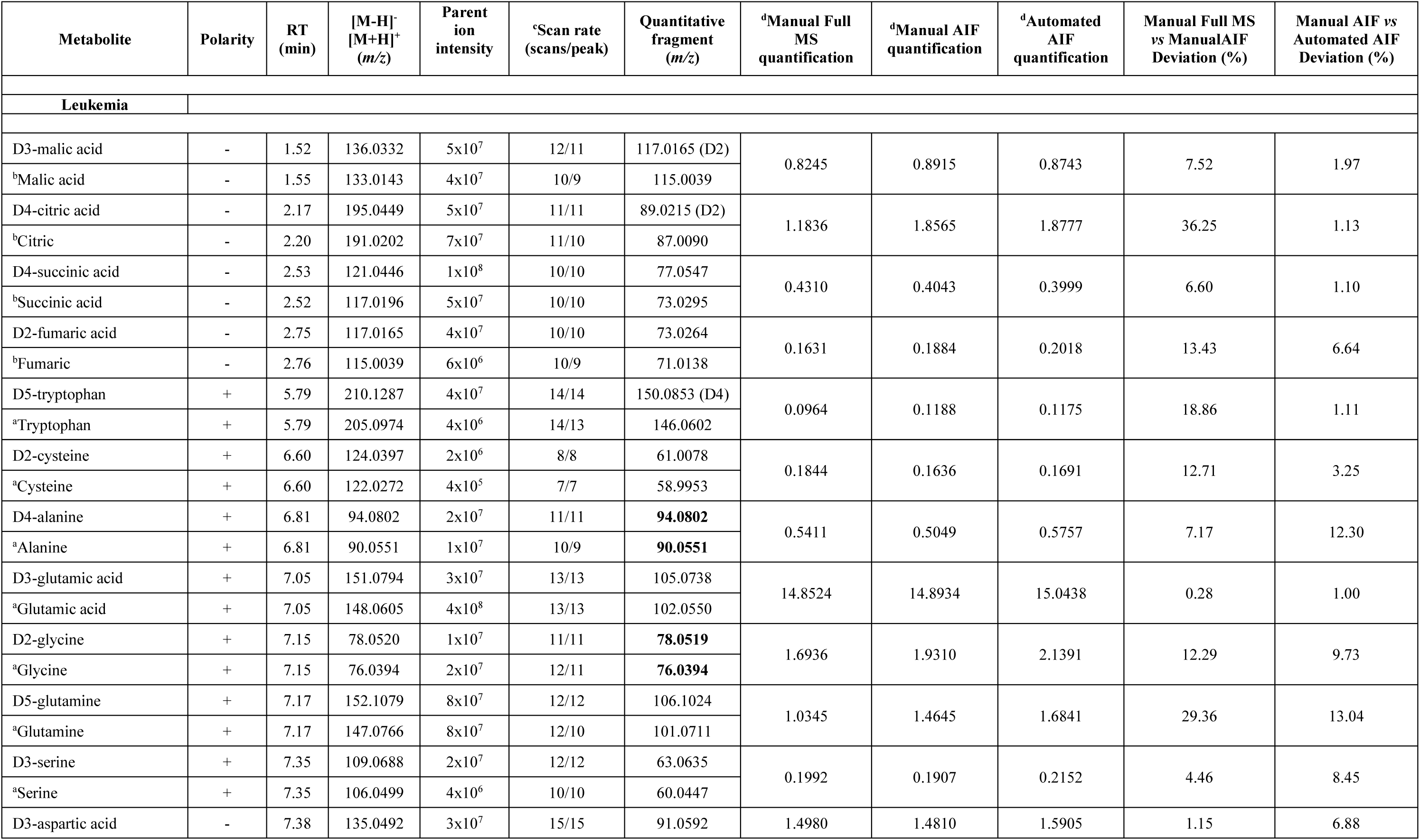
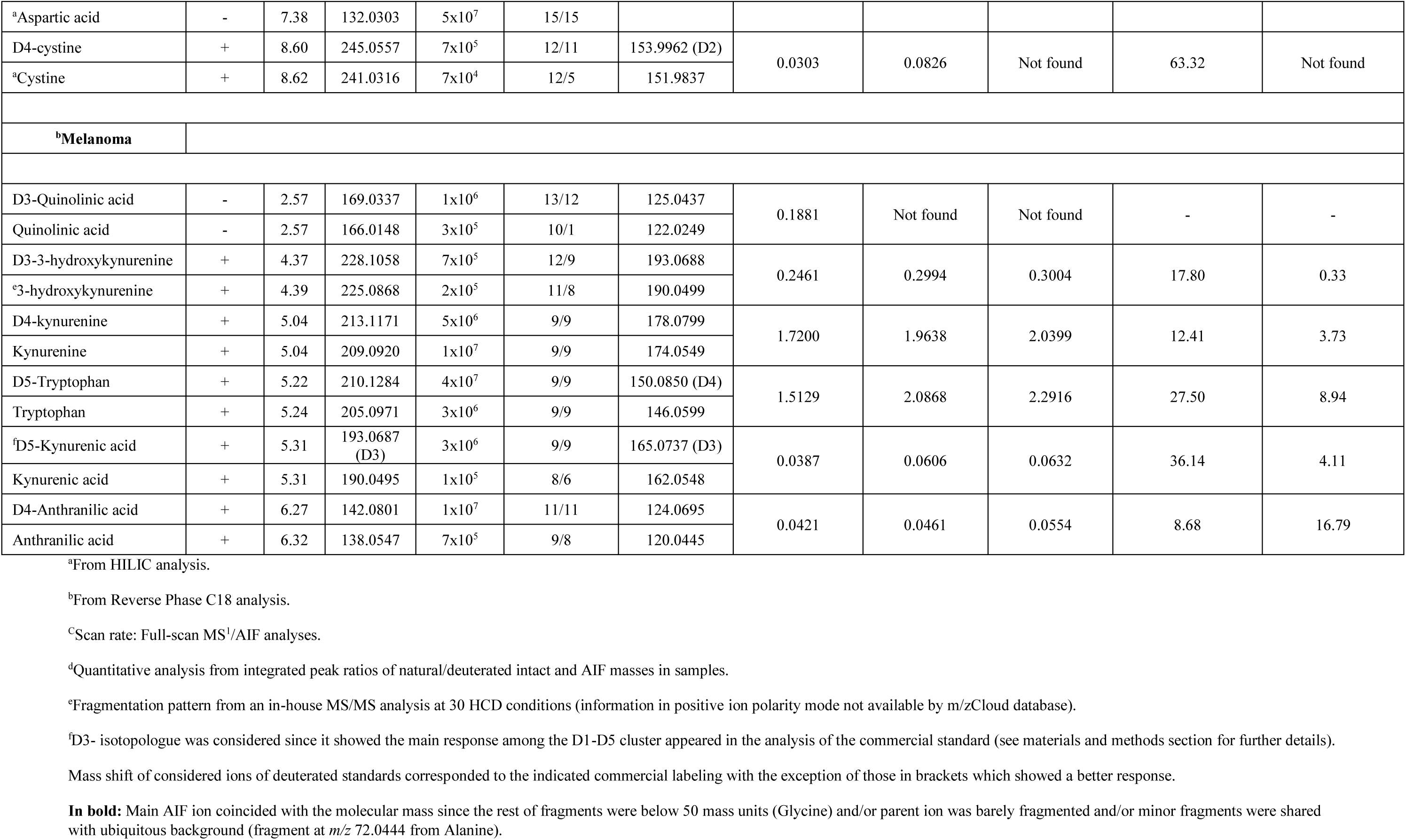
Metabolites determined in leukemia and melanoma samples by targeted full-scan MS^1^ and AIF analyses using deuterated standards.

Complementary information can also be extracted from scan rates, including changes in metabolite peak widths using different LC conditions. As an example, tryptophan had peak widths of 0.45 (14 scans/peak) and 0.3 (9 scans/peak) minutes with HILIC (Hydrophobic Interaction Liquid Chromatography) and reverse phase separation conditions, respectively. From **Table 1**, we can conclude that a minimum scan rate of 8 scans/peak is necessary to achieve reliable measurements in absence of interference from over fragmentation. Overall, peak representation of the AIF quantitative ions of deuterated standards exhibited high specificity and an absolute absence of artifacts from isobaric species from natural metabolites. This observation highlights the applicability of AIF analysis for isotope tracer studies, commonly used in metabolomics, due to the lack of interferences in fragments spectra [16].

### Targeted analysis: Manual vs. Automated AIF Analysis

Manually calculated AIF ratios were compared with those obtained from automated data processing done by R package MetaboList. Values from full MS ratios were not considered at this stage as only direct comparison of manual and automated AIF results was of interest, even for inaccurate quantifications (i.e. cystine) and poorly fragmented metabolites (alanine and glycine). As observed in Table 1, manual and automated AIF ratios achieved were comparable in almost all cases (deviations below 10%), thus providing evidence of the reliability of the automated data processing.

### Untargeted Analysis

Simultaneously to targeted analysis, automated AIF analysis enabled the untargeted qualitative metabolite profiling of samples. Results from MetaboList processing are shown in Tables S1 and S2 (see Supplemental Material). Each of which lists compounds in leukemia and melanoma samples, respectively, identified by their characteristic fragmentation patterns, beyond those studied using stable isotope-labeled standards. As a constraint, metabolite assignment required at least two characteristic fragments appropriately aligned with the respective parent ion peaks and signal-to-noise ratios above 10. Thus, identifications supported by only one fragment, as consequence of soft HCD-induced breakdown and/or below the limit of the considered mass range (m/z 50), such as lactic acid (m/z at 89.0244, same as parent ion) and cytidine (m/z at 112.0506, from the loss of ribose) were not included. As shown in **Tables S1** and **S2**, automated results from analysis with MetaboList are annotated (in the respective ionization mode) with the identified metabolites, their representative ions according to the MS level achieved (MS^1^ and MS^2^ for intact and AIF fragments, respectively), retention times of ions, peak widths, and integrated peak areas.

To test fragment alignment achieved by manual and automated processing, **Fig. 3** illustrates AIF results from Xcalibur and R-based processing of glucose found in leukemia sample considering its molecular mass at m/z 179.0563 and characteristic fragments at m/z 113.0245, 101.0245, 89.0246, 71.0137 and 59.0135 in negative ionization mode. Automated processing provided one single result (**Fig. 3G**) through the alignment of the intact mass and all five characteristic fragments in a time frame window of 6.48 – 6.51 minutes which was almost identical to results from manual analysis (**Figs. 3A**–**3F**). Appropriate alignment of masses represents an accurate alternative to the classical spectral matching procedure during the qualitative analysis of AIF data since relative abundances are poorly matched when comparing data-dependent breakdown patterns from precursors and AIF analysis.

**Figure 3.**
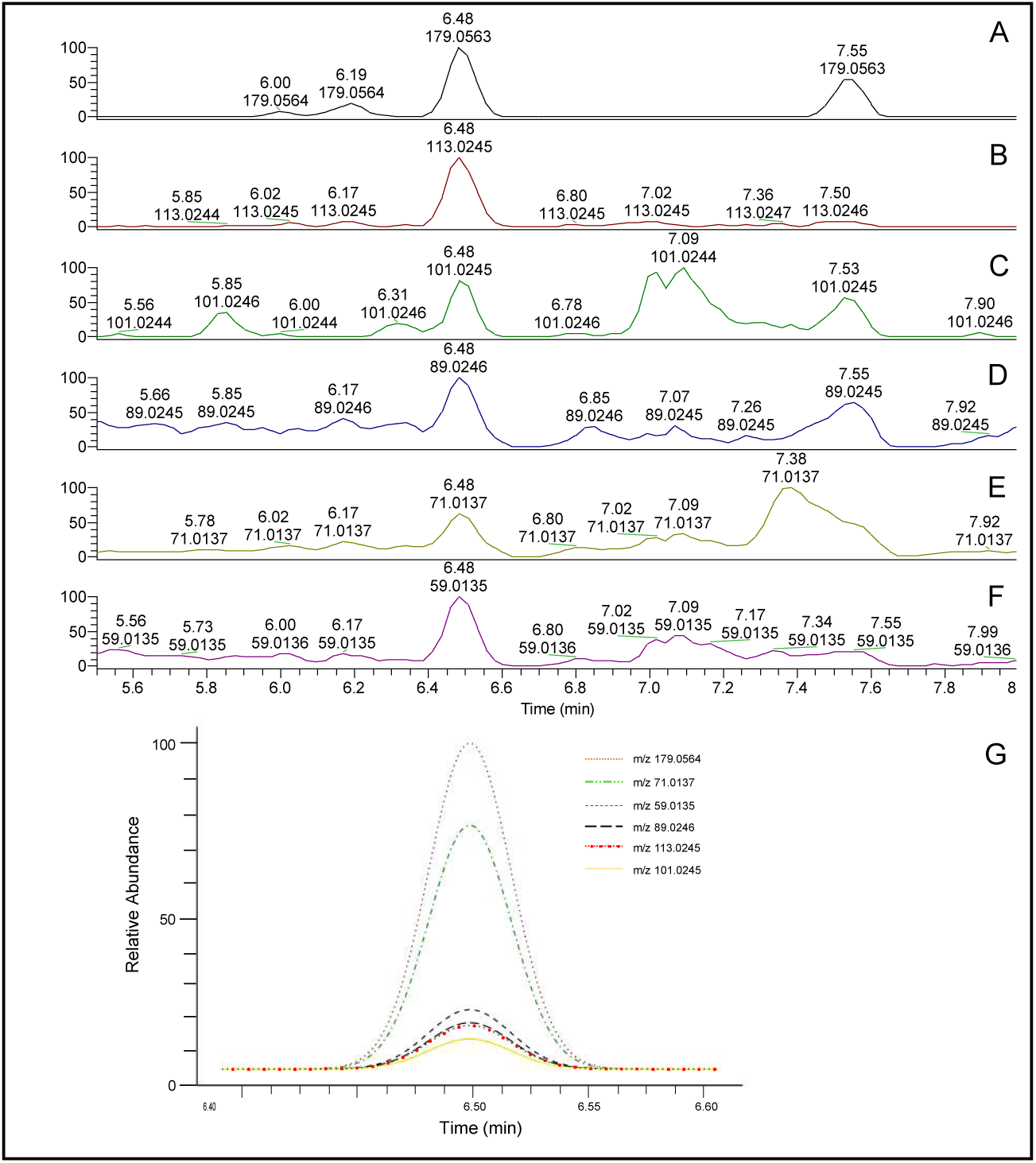
Manual alignment of characteristic ions of glucose (Table S1) from leukemia sample: A. Full MS-, B. AIF fragment at m/z-113.0245, C. AIF fragment at m/z-101.0245, D. AIF fragment at m/z-89.0246, E. AIF fragment at m/z-71.0137, F. AIF fragment at m/z-59.0135, G. Automated alignment of fragments.

Furthermore, the discrimination capacity of AIF analysis can be understood based on the results from leucine/isoleucine isomers found in leukemia sample in positive ionization mode. **Fig. 4A** shows the full MS representation of mass at *m/z* 132.1020 giving two peaks corresponding to the isomers. From the mzCloud database, leucine and isoleucine share two HCD ions at experimental *m/z* 132.1020 and 86.0965 (**Figs. 4B** and **4C**, respectively). Isomeric differentiation was achieved through the fragment at *m/z* 69.0700 that characterizes isoleucine (**Fig. 4D**). It should be noted that usefulness of the untargeted AIF analysis is restricted to metabolites that are accurately listed in the mzCloud database and commercially available standards from which it is possible to obtain characteristic high-resolution MS/MS fragmentation patterns. Furthermore, mzCloud dependency can be understood considering the extremely high mass accuracy provided by this available solution that greatly facilitates the building of reliable high-resolution in-house libraries, enabling users work within a 5 ppm mass tolerance range.

**Figure 4.**
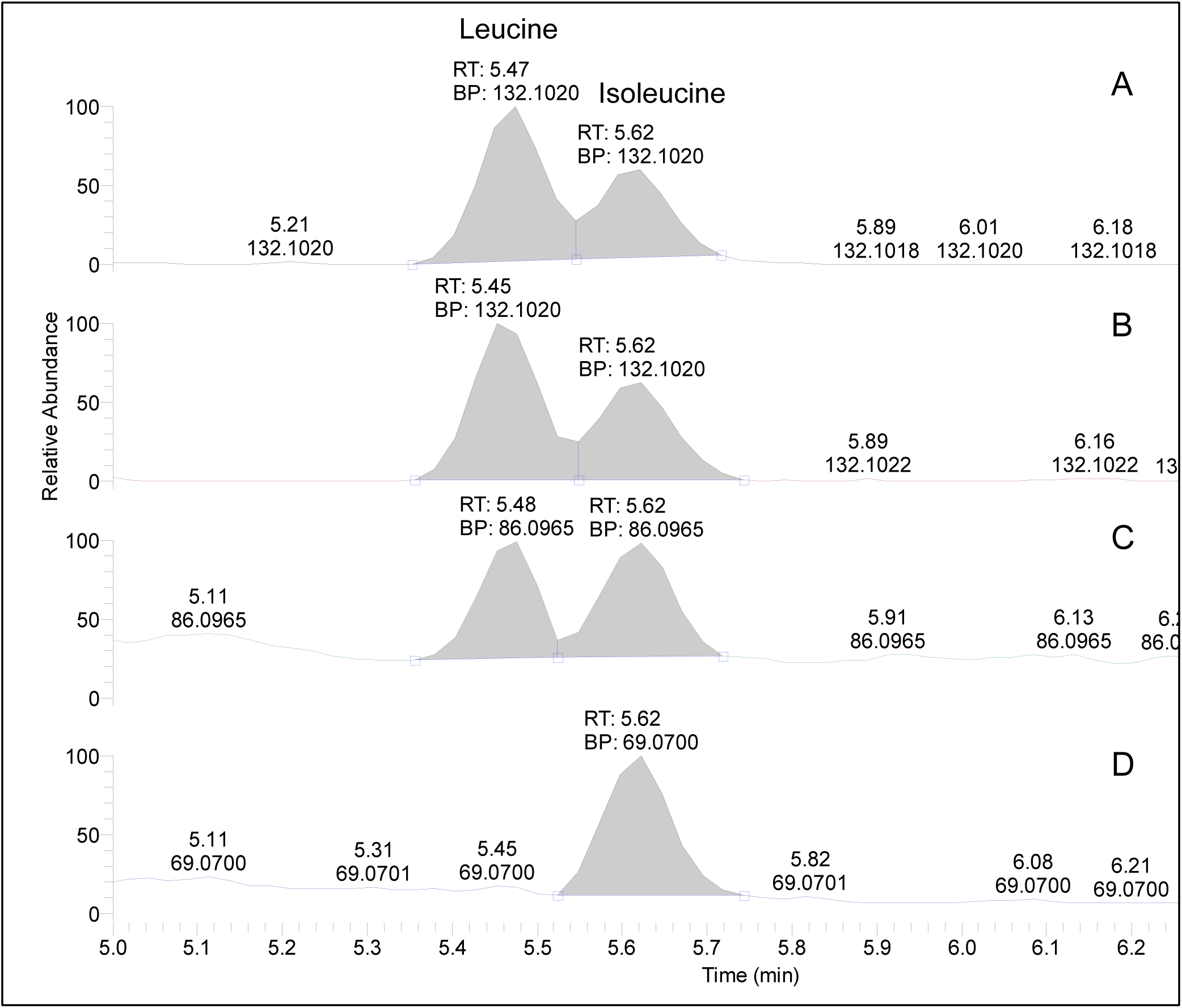
Full MS and AIF scan representation in positive ionization mode of leucine/isoleucine isomers in leukemia sample: A. Molecular mass at m/z 132.1020, B. AIF fragment at m/z 132.1020, C. AIF fragment at m/z 86.0965, D. AIF fragment at m/z 69.0700. Peak area integration from manual data processing indicated in gray. Nomenclature used: RT, chromatographic retention time; BP, base peak corresponding to intact and MS/MS ions (Table S1).

One of the advantages of using a highly efficient multiplexing MS system to clarify uncertainties is exemplified in **Table S2** by combining positive and negative ionization modes considering Adenosine-5-diphosphate (ADP) found in melanoma sample. Sensitivity of ADP is higher under positive ionization conditions however, qualitative analysis was performed in negative mode since six characteristic fragments (outside the intact mass) were identified in contrast to only one revealed in positive ionization mode. For the most complete untargeted analysis, multiplexing requires the use of fully flexible procedures to perform data processing, making the MetaboList package in R an ideal tool. Moreover, preliminary results (data not shown) demonstrated utility of R programming to perform background subtraction of AIF data to clarify the breakdown pattern of a considered metabolite. Thus, an isolated AIF spectrum of metabolites can be achieved by the subtraction of adjacent scans regarding the apex of their chromatographic peak, facilitating their positive assignment. Further studies are currently underway to refine our knowledge.

Untargeted qualitative analysis could likely be boosted by analyzing samples under saturating conditions to maximize the number of positive assignments. Saturating the detector with too many ions can increase the signal of poorly detected metabolites and their fragments. However, it should be taken into consideration that acquiring under saturating conditions sacrifices chromatographic peak shape, mass accuracy, and reproducibility, thus compromising the quality of quantitative analysis. In this research, samples were studied under non-saturating conditions to demonstrate the usefulness of the developed methodology to perform simultaneous quantitative/qualitative analyses. Furthermore, qualitative analysis can also be constrained by the level of exigence delimited by researchers when considering the number of representative breakdown fragments to be considered for positive assignments.

The methodology proposed in this research cannot be strictly considered as an untargeted approach at its initial stages of application. The untargeted analysis is limited by the number of metabolites listed in the in-house library built by users and from this, the extension of the aimed metabolite profiling will grow according to the continuous incorporation of positive assignments from samples and commercial standards. Once created, customized libraries are suitable to be implemented in the R package MetaboList workflow to carry out the analysis of samples independent of origin (biological fluids and tissues, foodstuffs, model solutions, natural extracts etc.). **Table S3** merges results achieved and can be used as an exportable MetaboList library. Thus, breakdown ions of positive assignments found in this research are listed in their appropriate polarity input format. Such exportable library is easily upgradable according to the desired goals of researchers worldwide.

## CONCLUSION

As demonstrated, implementation of AIF combined with R package MetaboList workflow analysis in metabolomics research was simple and comprehensive. No inclusion/exclusion precursor lists were required to obtain detailed intact and fragmented small (mostly *m/z* <250) polar metabolite profiles of leukemia and tumor samples assayed, thus avoiding inherent limitations of data-dependent MS/MS experiments. Qualitative and quantitative results were achieved by mimicking MRM analysis to simultaneously obtain targeted and untargeted results by combining full-scan MS and MS/MS analyses. Qualitative analysis resulted in high-confidence identification of small molecules, especially those with shared exact masses. Multiplexing capacity, outstanding mass accuracy, sensitivity, and relative error range of the orbitrap detector makes this device the ideal choice for merged scan modes. Efficiency of automated data processing of MS^1^ and AIF data through an affordable and entirely flexible open access R programming platform MetaboList was demonstrated. MetaboList may constitute an attractive alternative/complement to currently utilized solutions. This methodology is easily exportable and suitable to all MS devices with full-scan MS/MS capabilities independent of the manufacturer. Sensitivity is the main limitation of AIF analysis, but can be overcome by the increased scan speed of a new generation of analyzers. Future investigations should explore the expanding potential uses of AIF for targeted and untargeted metabolomics applications independent of sample nature.

## Acknowledgments

This work was supported by grants from the University of Texas System (ST: STAR Award), Hyundai Hope on Wheels research program, NIH R01 CA206210-01, and Cancer Prevention & Research Institute of Texas (CPRIT, DP150061).

